# FGF2-induced Redox Signaling: A Mechanism Regulating Pyruvate Dehydrogenase Driven Histone Acetylation and NANOG Upregulation

**DOI:** 10.1101/2023.01.20.524871

**Authors:** Petr Fojtík, Martin Senfluk, Katerina Holomkova, Anton Salykin, Jana Gregorova, Pavel Smak, Ondrej Pes, Jan Raska, Monika Stetkova, Petr Skladal, Miroslava Sedlackova, Ales Hampl, Dasa Bohaciakova, Stjepan Uldrijan, Vladimir Rotrekl

## Abstract

Precise control of pluripotency is a requirement for the safe and effective use of hPSCs in research and therapies. Here we report that pyruvate dehydrogenase upregulates histone H3 pan acetylation and levels of pluripotency marker NANOG in 5% O_2_. Pyruvate dehydrogenase (PDH) is an essential metabolic switch and a bottleneck for the glycolytic production of acetyl-CoA. Silencing of gene expression showed that PDH is regulated by the activity of its phosphatase PDP1. We show that PDP1 is sensitive to reactive oxygen species-mediated inactivation, leading to the downregulation of H3 pan acetylation and NANOG levels. Furthermore, we show that FGF2, a cytokine commonly used to maintain pluripotency activates pyruvate dehydrogenase through MEK1/2-ERK1/2 signaling pathway-mediated downregulation of ROS in 5% O_2_, thus promoting histone acetylation. Our results show the importance of pyruvate dehydrogenase in regulating energy metabolism and its connection to pluripotency. Furthermore, our data highlight the role of reactive oxygen species and redox homeostasis in pluripotency maintenance and differentiation.

**Highlights:**

- PDP1-induced activation of PDH leads to increased histone H3 pan acetylation and NANOG levels in hPSCs
- Reactive oxygen species (ROS) inactivate PDP1 and decrease histone H3 pan acetylation and NANOG levels in hPSCs
- MEK1/2-ERK1/2 signaling-mediated downregulation of ROS in 5% O_2_ activates PDH in hPSCs

**Graphical abstract:** 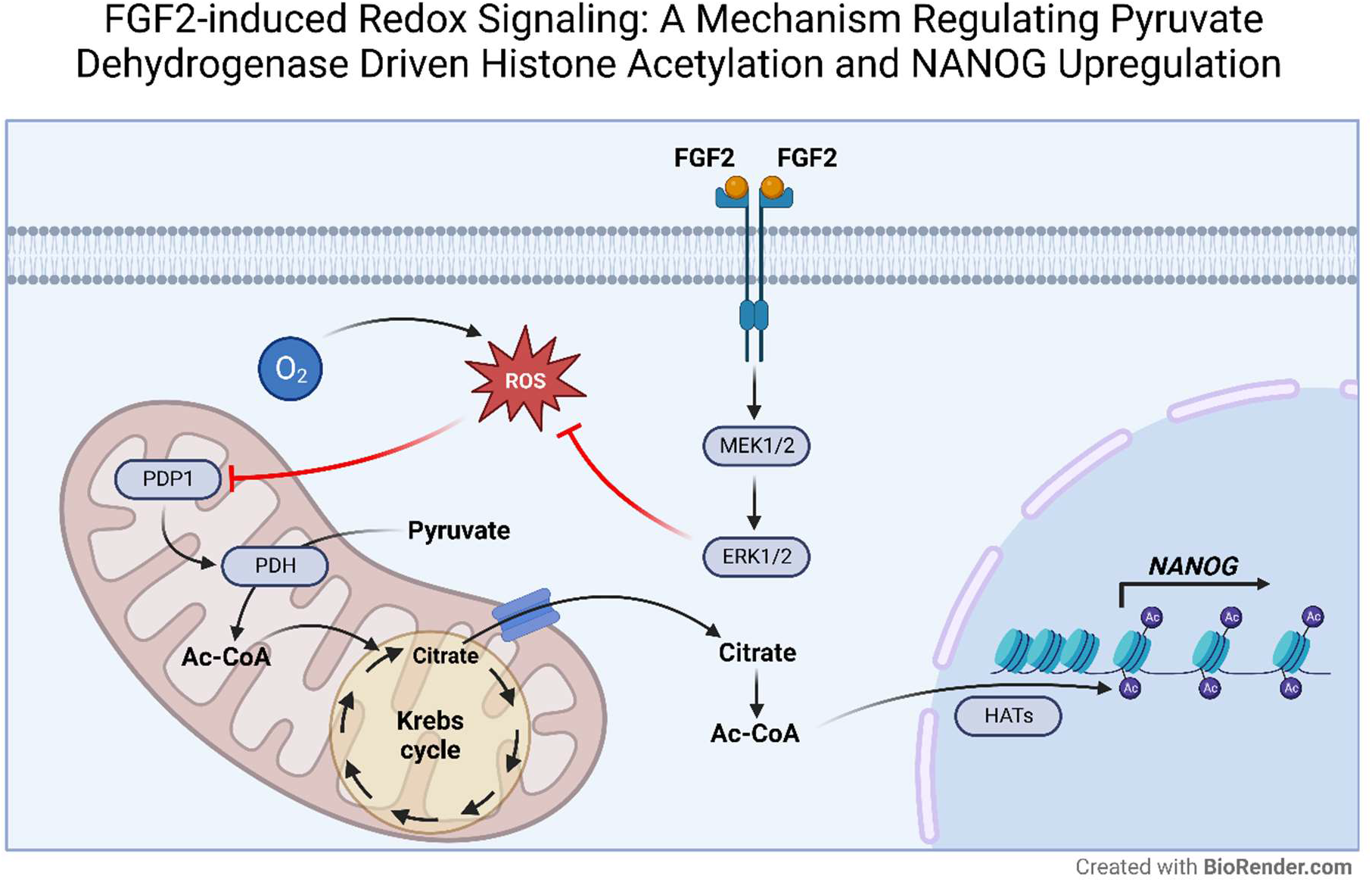

## Introduction

Human pluripotent stem cells (hPSCs) are an invaluable tool for developmental research, drug testing, and potentially for regenerative medicine. This is due to their ability to divide endlessly (self-renewal) and to differentiate into all cell types in our bodies (pluripotency). However, their safe and effective use depends on our ability to precisely control their fate determination, which proves difficult, as hPSCs show differentiation heterogeneity even among morphologically indistinguishable cellular populations (Kalmar et al., 2009; Nguyen et al., 2018; Hayashi et al., 2019). A significant part of the heterogeneity of cells, in general, stems from the differences in their epigenetic modifications (Carter and Zhao, 2020). Chromatin-opening histone modifications are upregulated in hPSCs to maintain their plasticity, with histone acetylation specifically recognized as one of the hallmarks of pluripotency (Mu et al., 2015; Huang et al., 2016; Trisciuoglio et al., 2018). To keep the histones acetylated, a supply of acetyl coenzyme A (Ac-CoA), a substrate for histone acetyltransferases (HATs), is required (Jo et al., 2020). Moussaieff and colleagues (Moussaieff et al., 2015) have shown that glycolysis is the primary source of Ac-CoA for histone acetylation in hPSCs. Concomitantly, hPSCs prefer glycolysis even when exposed to atmospheric oxygen, with only partial use of oxidative phosphorylation (OxPhos), as a source of glucose-derived energy (Varum et al., 2011; Tsogtbaatar et al., 2020).

Ac-CoA is produced in mitochondria from the end-product of glycolysis, pyruvate, by the pyruvate dehydrogenase complex (PDC). Ac-CoA can then continue in the tricarboxylic acid cycle (TCA) to fuel the OxPhos or be transported to the cytosol via a citrate shuttle. PDC is, therefore, an essential regulatory node in glucose metabolism linking glycolysis to Ac-CoA production and TCA. PDC consists of three enzymes (E1-E3), and its activity is mediated by the E1 subunit pyruvate dehydrogenase (PDH)(Patel and Korotchkina, 2001). It can be deactivated by pyruvate dehydrogenase kinases (PDHKs, isoforms 1-4) induced phosphorylation and activated by dephosphorylation by pyruvate dehydrogenase phosphatases (PDPs, isoforms 1-2) (Patel and Korotchkina, 2006). As glycolysis is the primary source of Ac-CoA for histone acetylation in hPSCs (Moussaieff et al., 2015), we hypothesized that modulation of PDH activity would have a significant impact on histone acetylation and pluripotency. Furthermore, PDP1 supports histone acetylation in melanoma cells (Karagiota et al., 2022). To uncover the mechanisms regulating PDH, we explored the effects of fibroblast growth factor 2 (FGF2) signaling and low oxygen tensions on PDH phosphorylation, as both are known for upregulation of pluripotency (Eiselleova et al., 2009; Mathieu et al., 2013; Närvä et al., 2013; Haghighi et al., 2018) and glycolysis (Papandreou et al., 2006; Fumarola et al., 2017; Liu et al., 2018).

In vitro pluripotency maintenance of hPSCs depends on downstream signaling of several cytokines like FGF2 (Eiselleova et al., 2009). FGF2 induces pluripotency genes through activation of its downstream MEK1/2-ERK1/2 (Li et al., 2007; Haghighi et al., 2018) and PI3K/AKT (Wang et al., 2017) signaling pathways. Apart from the direct effect on crucial pluripotency transcription factors like OCT-4, SOX2, or NANOG, these pathways were also shown to support glycolytic metabolism favored by hPSCs. ERK1/2 was found to upregulate GLUT1 glucose transporter, lactate dehydrogenase, and pyruvate kinase M2 (Yang et al., 2012). The PI3K/AKT pathway and its downstream effector mTOR govern anabolic reactions, nutrient uptake, cell growth, and survival (Robey and Hay, 2009; Saxton and Sabatini, 2017). PI3K/AKT is also an essential regulator of aerobic glycolysis, as it increases glucose uptake by upregulation of GLUT1 (Zhou et al., 2020), hexokinase (Gottlob et al., 2001), and phosphofructokinase 1 kinase (Deprez et al., 1997).

Reduced oxygen tensions (5% O_2_) are beneficial for embryo development (Dunwoodie, 2009) as well as for hPSCs culture as they prevent differentiation (Lin et al., 2006; Närvä et al., 2013) and enhance the generation of hiPSCs (Yoshida et al., 2009). Furthermore, the transition of hPSCs committed to differentiation to a low O_2_ environment led to a reversion of the differentiation process (Mathieu et al., 2013). Low oxygen tensions also lead to the utilization of glycolysis with limited OxPhos favored by hPSCs (Kim et al., 2006). The main facilitators of cellular adaptation to low oxygen tensions are hypoxia-inducible factors (HIFs) (Semenza, 2001). HIFs-upregulated expression of PDHKs leads to inhibition of PDH, ultimately decreasing the flow of metabolites to the Krebs cycle, and consequently downregulates OxPhos (Kim et al., 2006). In addition, to compensate for the reduction in energy production, HIFs upregulate the enzymes of glycolysis (Semenza et al., 1996).

Low oxygen tensions and stabilization of HIFs also help to downregulate reactive oxygen species (Samanta and Semenza, 2017; Fojtík et al., 2021), which were shown to modulate various cellular processes like signaling pathways (Okoh et al., 2013; Zhang et al., 2016; Fojtík et al., 2021) or energy metabolism through inhibition of PDH (Tabatabaie et al., 1996; Humphries and Szweda, 1998). These effects are facilitated through the reversible oxidation of various enzymes like protein phosphatases (Sommer et al., 2002; Östman et al., 2011), including PDH phosphatases (Wright et al., 2009). As we have shown that ROS also downregulate pluripotency markers (Fojtík et al., 2021), we further hypothesized that this might be due to the inhibition of PDH and downregulation of histone acetylation.

In this work, we demonstrate that PDH serves as a vital gatekeeper of histone acetylation in hPSCs that is regulated by intracellular ROS levels, which are, in turn, controlled by FGF2-MEK1/2-ERK1/2 signaling and oxygen levels. Furthermore, we show that the protein levels and expression of the key pluripotency gene NANOG depend on the PDH-induced production of Ac-CoA. Finally, we identify oxidative inhibition of PDP1 as the mechanism behind ROS-induced downregulation of PDH activity.

## Results

Glycolysis is the primary source of Ac-CoA for histone acetylation (Moussaieff et al., 2015), a major pluripotency marker of hPSCs (Mu et al., 2015; Qiao et al., 2015; Huang et al., 2016; Trisciuoglio et al., 2018). We hypothesized that pyruvate dehydrogenase (PDH), an enzyme connecting glycolysis to TCA and Ac-CoA production, might be regulated by FGF2 signaling, as FGF2 is involved in the control of glycolysis (Fumarola et al., 2017; Liu et al., 2018). Furthermore, we investigated the effect of FGF2 on PDH at 21% and 5% O_2_, as low O_2_ has been shown to promote pluripotency (Lin et al., 2006; Yoshida et al., 2009; Forristal et al., 2010; Mathieu et al., 2013), modulate energy metabolism (Semenza, 2011), and we described distinct changes in FGF2 downstream signaling in response to oxygen levels (Fojtík et al., 2021).

### FGF2-induced activation of PDH is amplified in low O_2_ and independent of PDHK1 levels

To analyze the regulation of PDH activity, we studied the protein amount of pyruvate dehydrogenase kinase 1 (PDHK1), pyruvate dehydrogenase phosphatase 1 (PDP1), total levels of PDH, and PDH phosphorylation on Ser^293^, which is the most abundantly phosphorylated site on PDH (Rardin et al., 2009), using Western blot (WB) (Fig. 1A). To investigate the effect of FGF2 in different oxygen concentrations, hPSCs were treated with FGF2 (10ng/ml) under a 21% and 5% O_2_ atmosphere. FGF2-starved (for 24 h) hPSCs were treated with FGF2 for 24 hours and then for another hour after media change to promote the FGF2 signaling since FGF2 has low thermostability (Chen et al., 2012). This way, we were able to capture the immediate effects of FGF2 signaling, as well as those that might require more time to manifest. FGF2-untreated cells were used as a negative control. In 5% O_2_, FGF2 significantly decreased phosphorylated PDH and, at the same time, significantly increased the amount of PDHK1 compared to FGF2-untreated cells, while levels of PDP1 remained unchanged (Fig. 1A). A similar effect of FGF2 on PDHK1 and PDP1 levels was observed in 21% O_2_, but the change in PDH phosphorylation was not statistically significant. The overall phosphorylation levels of PDH in 5% O_2_ were significantly higher compared to the corresponding treatment in 21% O_2_.

**Figure 1.**
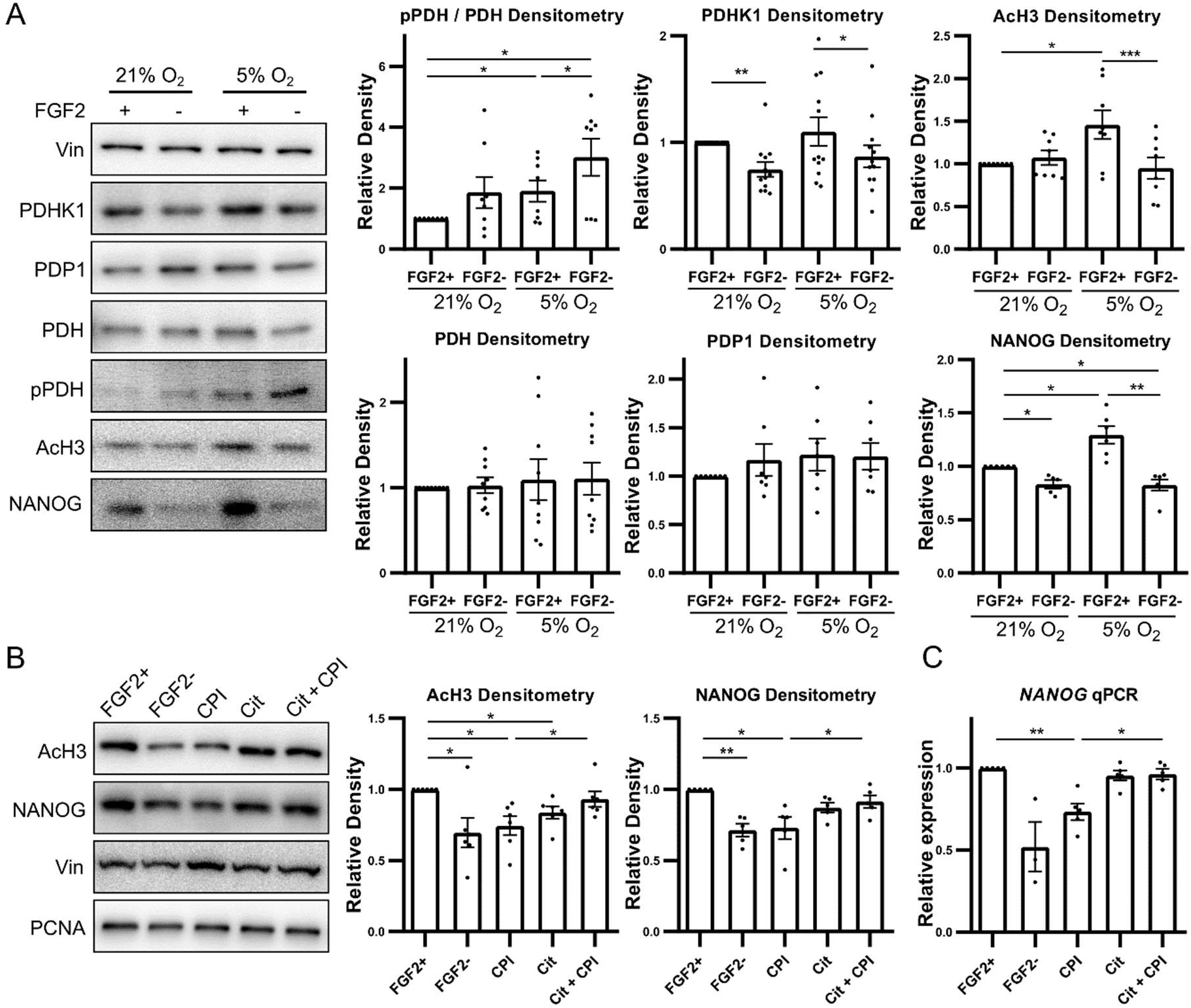
FGF2 activates pyruvate dehydrogenase in 5% O_2_ leading to histone acetylation. **A)** Western blot and densitometric analysis of PDH (N=9), PDHK1 (N=12), PDP1 (N=7), NANOG (N=6), H3 pan-acetylation (AcH3; N=8) and a ratio of phosphorylated to total PDH (N=8). A significant decrease in PDH phosphorylation was accompanied by a substantial increase in PDHK1, AcH3, and NANOG amount in FGF2 treated cells in 5% O_2_. Vinculin (Vin) was used as a loading control. **B)** WB and densitometric analysis of H3 pan-acetylation (AcH3; N=6) and NANOG (N=5) amount after PDH inhibition and rescue of the effect by sodium citrate (Cit). CPI treatment led to a significant downregulation of AcH3 and NANOG comparable to the negative control, which was rescued by sodium citrate. Vinculin (Vin) and PCNA were used as loading controls. **C)** Analysis of *NANOG* expression (N=5) using qPCR after PDH inhibition and rescue of the effect by sodium citrate (Cit). Treatment with CPI led to a significant decrease in *NANOG* expression, which was rescued by the addition of Cit.

This enzymatic machinery resides in mitochondria. Mitochondria in hPSCs are localized perinuclearly, are small, and with undeveloped cristae (Choi et al., 2015). To determine the potential effect of FGF2 on mitochondrial morphology, we performed staining using a MitoTracker Red CMXRos probe and transmission electron microscopy of mitochondria in cells treated with and starved of FGF2 for 24 hours in 21% O_2_. We did not observe changes in mitochondrial size or development (Figure S1). After inhibiting PDHK1 with dichloroacetate (DCA; 20mM/24h), we observed the elongation of mitochondria and development of their cristae (Figure S1), with no effect of FGF2 on this process.

### FGF2 promotes pluripotency in a low oxygen-dependent manner by global histone acetylation mediated by PDH activation

To assess the changes in histone acetylation, we inspected the pan-acetylation of histone H3 (AcH3) using WB and an antibody specific for all five acetylation sites on H3 (acetyl K9 + K14 + K18 + K23 + K27). Indeed, the cells treated with FGF2 exhibited significantly higher levels of AcH3 in 5% O_2_ (Fig. 1A), suggesting that FGF2 treatment promotes histone acetylation in a low-oxygen environment. However, FGF2 did not increase the levels of pan acetylation of H3 in 21% O_2_. To elucidate the impact of the FGF2-mediated increase in AcH3 on pluripotency, we looked at the levels of pluripotency marker NANOG using WB. FGF2 treatment significantly increased the levels of pluripotency marker NANOG regardless of O_2_ concentration, but NANOG levels were significantly higher in 5% O_2_ FGF2-treated cells compared to FGF2-treated cells in 21% O_2_ (Fig. 1A).

To clarify whether the increase in histone acetylation and NANOG is caused by FGF2-mediated activation of PDH in 5% O_2_, we inhibited PDH using a small molecule inhibitor CPI-613 (CPI; 10μM) (Smith and Hewitson, 2020) for 24 hours and additional 2 hours after media change. The data show that the increase in PDH phosphorylation in FGF2-starved cells or inhibition of PDH by CPI for 24 hours similarly result in significantly lower levels of AcH3 (Fig. 1B), which is accompanied by a significant decrease in protein levels and expression of the pluripotency marker NANOG (Fig. 1B, C). This correlation suggests that FGF2-induced PDH activation is needed for histone acetylation in hPSCs in 5% O_2_ and might impact pluripotency. To test whether the PDH-produced Ac-CoA is indeed required for histone acetylation and NANOG production, we rescued the inhibition of PDH by CPI using 5mM sodium citrate, which bypassed the production of Ac-CoA by PDH (Petillo et al., 2020) (Fig. 1D). Treatment with sodium citrate significantly increased levels of pan acetylation of H3 and NANOG levels in CPI treated cells rescuing the effect of PDH inhibition, while it did not exhibit any significant effect by itself (Fig. 1. B, C). This data confirms that FGF2-activated PDH is a crucial gatekeeper to produce Ac-CoA used in histone acetylation and that NANOG levels depend on this mechanism in hPSCs cultured at 5% O_2_.

### Reactive oxygen species modulate PDH activity by inhibiting PDP1

We have previously reported that 5% O_2_ decreases levels of the crucial second messengers — reactive oxygen species (ROS) — enough to modulate FGF2 signaling, and that FGF2 signaling itself likely downregulates ROS as MAPK inhibition led to upregulation of ROS in 5% O_2_ (Fojtík et al., 2021). This was confirmed as we observed that FGF2 downregulates ROS in 5% O_2_ (Fig. 2A) in cells starved of (FGF2-) or treated with FGF2 (FGF2+; 10 ng/ml) for 24 hours and an additional 2 hours after media change. The FGF2-induced significant increase of histone H3 pan acetylation observed exclusively in a 5% O_2_ environment (Fig. 1A) suggests this mechanism is oxygen-sensitive. To analyze the involvement of ROS signaling in PDH regulation, we induced ROS in hPSCs cultivated at 5% O_2_ by adding 5μM H_2_O_2_ for 24 hours and additional 2 hours after media change. H_2_O_2_-treated cells exhibited a significant rise in PDH phosphorylation, accompanied by a substantial decrease in AcH3 and NANOG levels (Fig. 2B).

**Figure 2.**
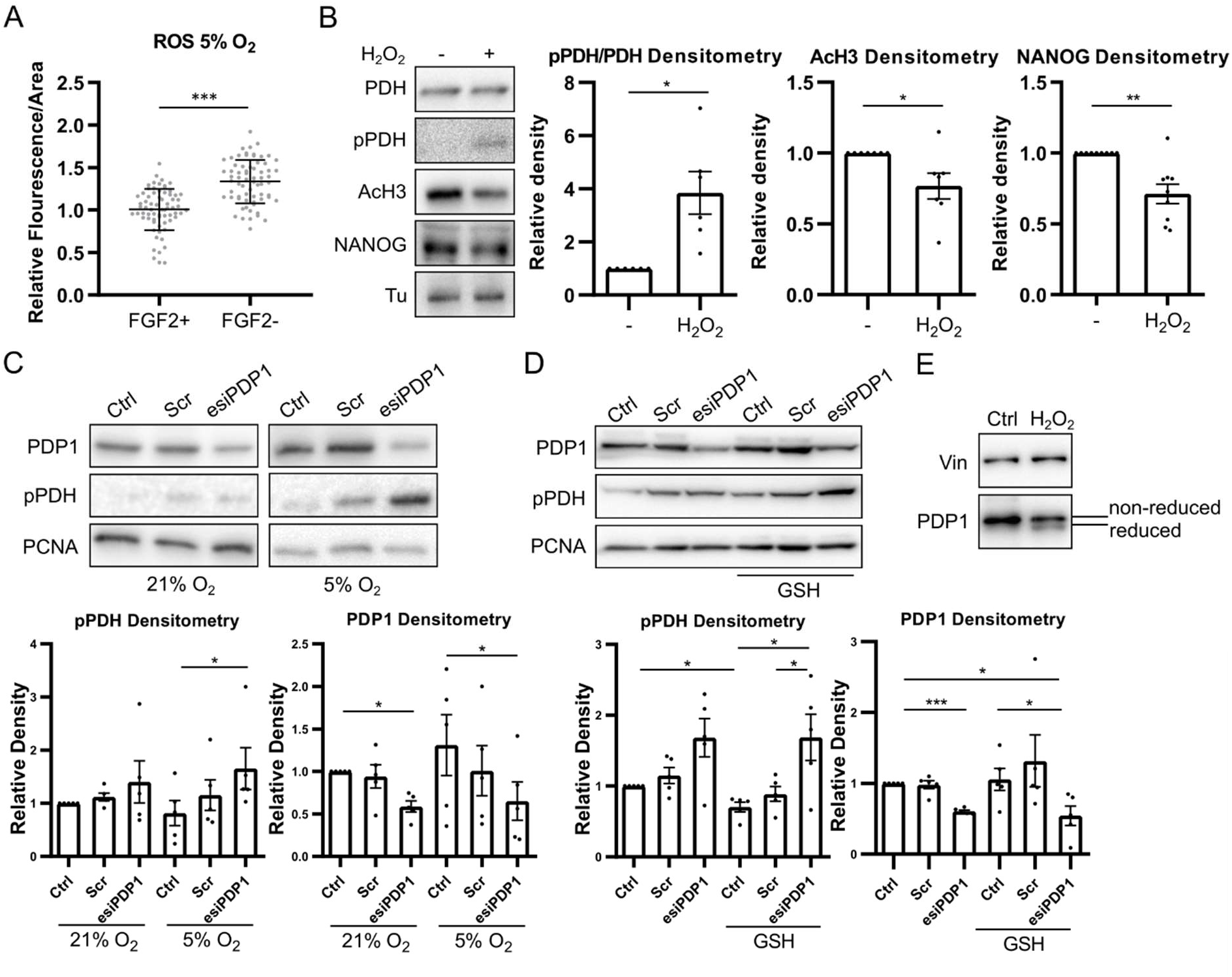
FGF2-activated PDH leads to the utilization of the Krebs cycle in 5% O_2_. **A)** Quantification of ROS measurement using CellROX Green in 5% O_2_ in CCTL14 hPSCs. Normalized data from 5 different experiments (with at least 10 measurements in each) are presented as mean ± SD (FGF2+ N=70; FGF2-N=67). Statistical significance was calculated using an unpaired two-tailed t-test. **B)** ROS increase PDH phosphorylation (pPDH/PDH N=6) and decrease AcH3 (N=7) and NANOG (N=9) levels. WB analysis and densitometry of samples from CCTL14 hPSCs. Tubulin (Tu) was used as a loading control. **C)** Silencing of PDP1 expression increases PDH phosphorylation in 5% O_2_. WB analysis and densitometry of pPDH (N=5) and PDP1 (N=5) from CCTL14 hPSCs. Cells transfected with scramble siRNA were used as a negative control. PCNA was used as a loading control. **D)** Silencing of PDP1 expression increases PDH phosphorylation in GSH-treated cells in 21% O_2_. WB analysis and densitometry of pPDH (N=5) and PDP1 (N=5) from CCTL14 and CCTL12 hPSCs. Cells transfected with scramble siRNA were used as a negative control. PCNA was used as a loading control. **E.** WB analysis of PDP1 oxidation on CCTL14 hPSCs. Oxidized PDP1 migrated faster on the 8% gel.

Next, we looked at the possibility of ROS-mediated regulation of PDP1 as phosphatases are prone to be regulated by ROS (Sommer et al., 2002; Wright et al., 2009), and PDP1 was observed to regulate histone acetylation (Karagiota et al., 2022). Because we observed a stronger effect of FGF2 on PDH phosphorylation in low ROS 5% O_2_ compared to 21% O_2_ (Fig. 1A), we first analyzed the difference in PDP1 activity in hPSCs in 21% and 5% O_2_. To gauge the PDP1 activity, we silenced its expression using small interfering RNA and compared the change in phosphorylation of its target PDH with the control. The results show that PDP1 silencing significantly increased PDH phosphorylation at 5% O_2_ while having only a partial effect in 21% O_2_ (Fig. 2C). To confirm the involvement of higher ROS levels in 21% O_2_ in PDP1 inactivation, we treated the PDP1-silenced cells in 21% O_2_ with reduced glutathione (GSH; 5mM/1h). Indeed, we observed a statistically significant increase in PDH phosphorylation in GSH-treated cells with silenced PDP1 compared with the corresponding control (Fig. 2D), further suggesting that the endogenous ROS levels at 21% O_2_ are sufficient for PDP1 reversible oxidation and inhibition. We also observed a decreased PDH phosphorylation in GSH-treated controls compared to untreated control hPSCs. Furthermore, WB analysis of PDP1 in non-reducing conditions revealed a mobility shift in H_2_O_2_-treated cells, suggesting the possibility of oxidation of PDP1 (Fig. 2E).

### FGF2 regulates PDH activity by MEK1/2-ERK1/2 mediated downregulation of ROS

We have previously reported that FGF2-activated MEK1/2-ERK1/2 signaling downregulates ROS levels in hPSCs (Fojtík et al., 2021). Here we decided to test whether ROS-increasing MAPK inhibition influences PDH phosphorylation described in melanoma cells (Cesi et al., 2017) and, subsequently, acetylation of histones. We treated hPSCs with a specific MEK inhibitor PD03 (PD0325901) (0.2μM/2 h) in 21% and 5% O_2_ atmosphere and observed a distinct effect based on the O_2_ concentration (Fig. 3A). MEK1/2 inhibition in 21% O_2_ did not significantly increase PDH phosphorylation. In contrast, MEK1/2 inhibition in 5% O_2_ significantly increased PDH phosphorylation to levels comparable with FGF2-starved cells. We did not observe an effect on PDHK1 levels. The increase in PDH phosphorylation was accompanied by a significant decrease in mitochondrial membrane potential in 5% O_2_ after MEK1/2 inhibition (Fig. 3B). To test if the upregulation of ROS is responsible for the increase in PDH phosphorylation after MEK1/2 inhibition, we combined the PD03 (0.2μM/1h) treatment with GSH (5mM/1h), which quenched ROS and decreased the PD03-induced PDH phosphorylation (Fig. 3C), suggesting that MEK1/2-ERK1/2 downregulates PDH phosphorylation, at least partially, via ROS quenching. To decouple the ROS effect on PDH phosphorylation from the increased phosphorylation of AKT observed after MEK1/2-ERK1/2 inhibition upon PD03 treatment (Fojtík et al., 2021), we treated FGF2-starved cells (24h) cultivated in 5% O_2_ with an AKT activator SC79 (10μM/2h) (Zhu et al., 2019) and FGF2 (10 ng/ml/2h). The treatment with SC79 led to an increase in AKT phosphorylation in 5% O_2_, but we did not observe any effect on PDH phosphorylation (Fig. 3D).

**Figure 3.**
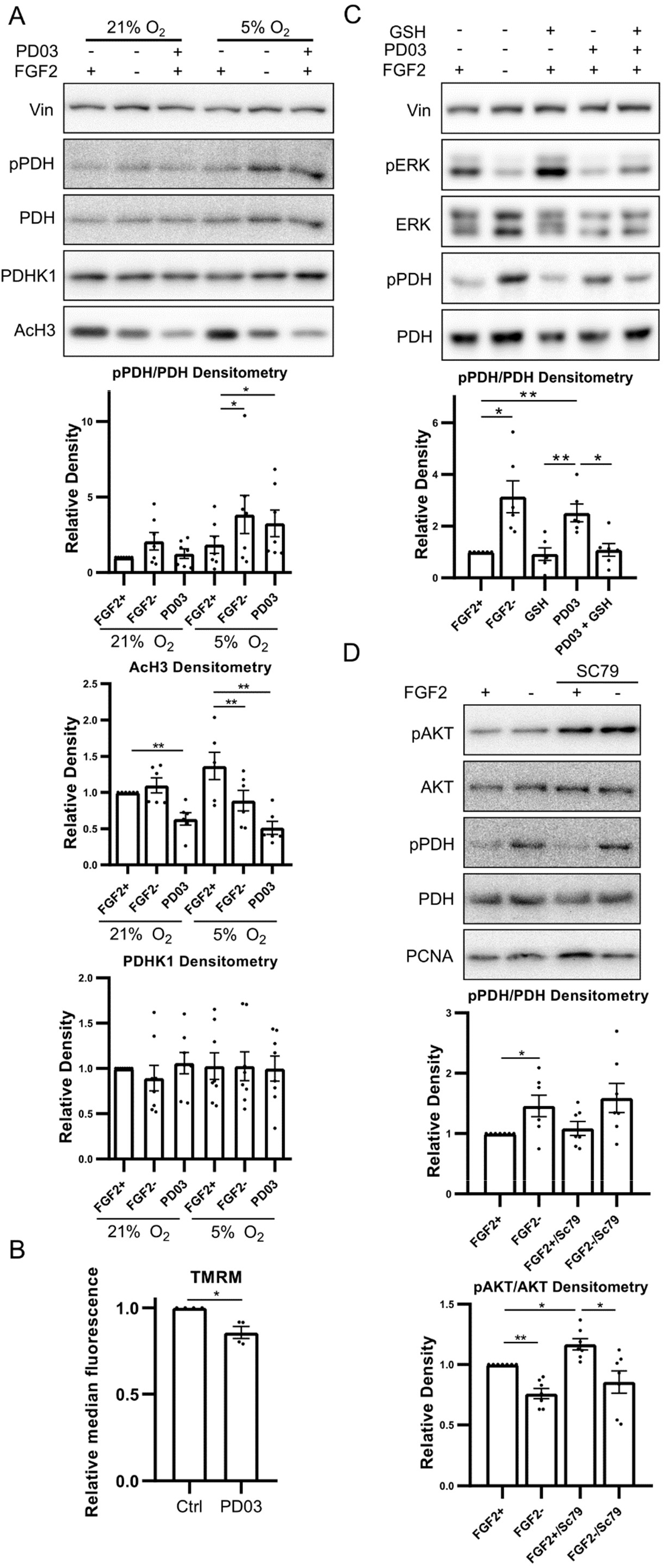
PDH phosphorylation is downregulated by MEK1/2-ERK1/2-mediated downregulation of ROS. **A)** WB analysis and densitometry of the effect of PD03-induced MEK1/2 inhibition on PDH phosphorylation in CCTL14 and AM13 hPSCs. WB analysis and densitometry of pPDH/PDH (N=7), PDHK1 (N=8), and AcH3 (N=6) MEK1/2 inhibition led to a significant decrease in AcH3 regardless of the O_2_ concentration and pPDH upregulation exclusively in 5% O_2_. Vinculin (Vin) was used as a loading control. **B)** Measurement of mitochondrial membrane potential after MEK1/2 inhibition in 5% O_2_ in CCTL14 hPSCs. Data represent the normalized median fluorescence ± SEM (N=4). Statistical analysis was performed using one sample t-test (theoretical mean = 1). **C)** WB analysis of GSH-mediated rescue of MEK1/2 inhibition-induced PDH phosphorylation in CCTL14 hPSCs cultivated in 5% O_2_ (N=6). GSH rescued the MEK1/2 inhibition-induced pPDH upregulation. PCNA was used as a loading control. **D)** WB analysis of the effect of AKT activation on PDH phosphorylation in 5% O_2_. SC79 treatment induced AKT phosphorylation but did not affect the proportion of phosphorylated PDH (N=7). PCNA was used as a loading control.

To further verify the activity of unphosphorylated PDH in FGF2-treated hPSCs in 5% O_2_, we measured mitochondrial transmembrane potential that reflects the activity of the electron transport chain, oxygen consumption rates, and concentration of several metabolites of the Krebs cycle. For these experiments, hPSCs were treated with FGF2 (10 ng/ml) for 24 hours and again the following day for 2 hours; the data were then compared to FGF2-starved cells. Our data show a small but statistically significant increase in mitochondrial transmembrane potential in FGF2-treated cells (Fig. 4A). Oxygen consumption rate, measured using a Clark-type electrode, correlates with the increased mitochondrial potential as it is significantly higher in FGF2-treated cells in 5% O_2_ (Fig. 4B). Both results are in agreement with the higher activity of PDH in FGF2-treated cells.

**Figure 4.**
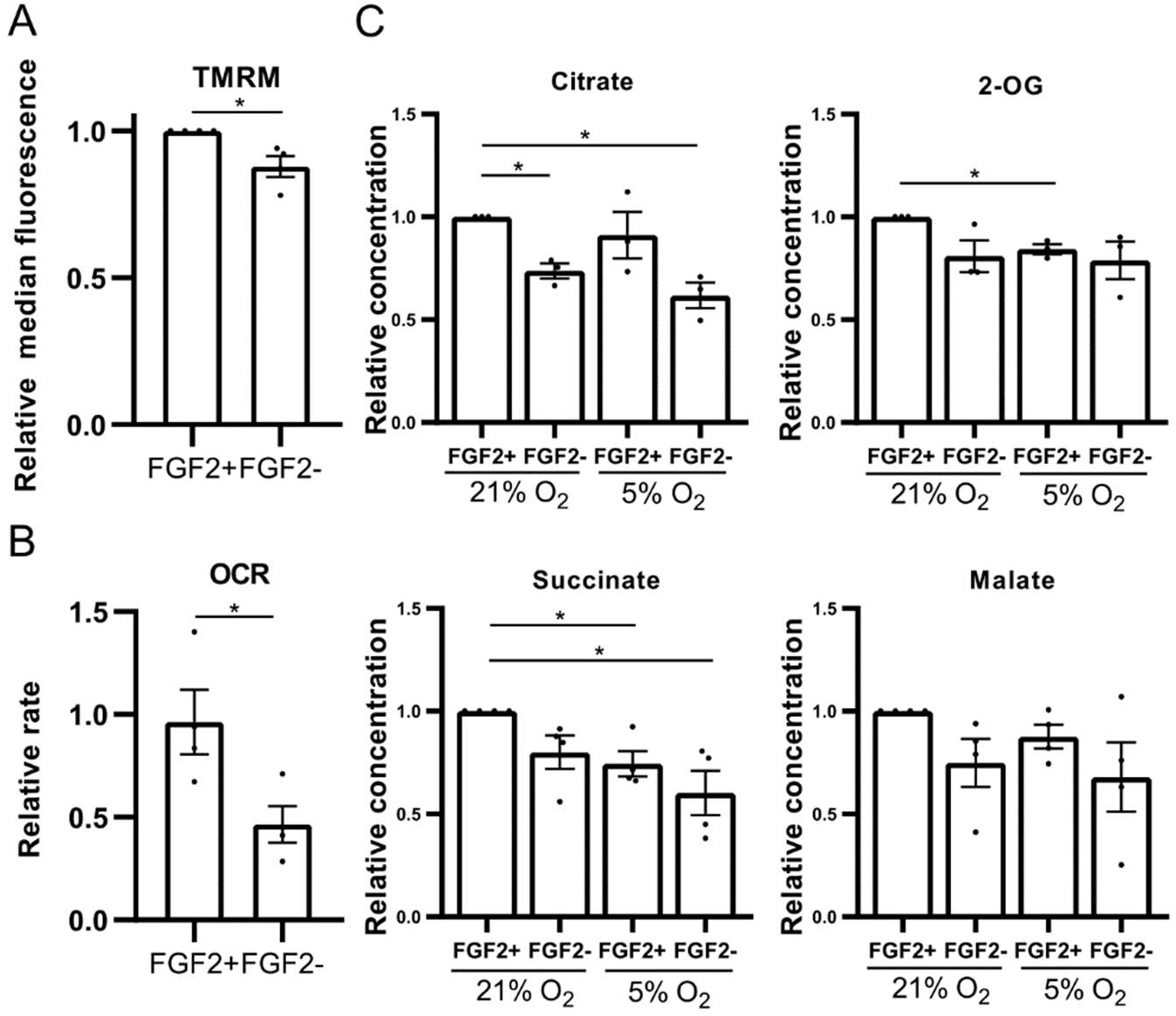
FGF2 leads to the utilization of OxPhos in 5% O_2_. **A)** Measurement of the effect of FGF2 on mitochondrial membrane potential in 5% O_2_ in CCTL14 hPSCs. Data represent the normalized median fluorescence ± SEM (N=4). **B)** Measurement of oxygen consumption rates (OCR) in 5% O_2_ in CCTL14 hPSCs. FGF2-treated cells show significantly increased oxygen consumption rates. Data represent the relativized OCR (nM/million cells/min) ± SEM (N=4). Statistical analysis was conducted using a two-tailed unequal variances t-test. **C)** Measurement of selected Krebs cycle metabolites content in CCTL14 hPSCs cultivated in 21% and 5% O_2_. Relativized concentration of citrate (N=3), 2-oxoglutarate (2-OG; N=3), succinate (N=4), and malate (N=4).

Using liquid chromatography coupled with mass spectrometry, we determined the concentration of products of several steps of the TCA cycle in cells starved of FGF2 for 24 hours and then treated with FGF2 (10 ng/ml) for 24 hours and additional 2 hours after media change. The data show that in 5% O_2_, the amount of citrate in FGF2-treated cells was comparable to FGF2-treated cells in 21% O_2_, corresponding to lower PDH phosphorylation (higher activity) (Fig. 4C). Interestingly, the amount of products of the subsequent steps of TCA, 2-OG and succinate, has significantly decreased in 5% O_2_ FGF2-treated cells compared to the same treatment in 21% O_2_. Furthermore, we observed a statistically significant increase in citrate in FGF2 -treated cells in 21% O_2_ compared to the untreated control and a non-significant increase in 2-OG, succinate, and malate (Fig. 4C). Taken together, our data suggest that PDH is indeed more active in FGF2-treated hPSCs, which correlates with our observations of PDH phosphorylation (Fig. 1A).

## Discussion

FGF2-induced MEK1/2-ERK1/2 signaling is instrumental for hPSCs in maintaining their pluripotency (Haghighi et al., 2018). Also, glucose metabolism has been increasingly recognized as an important part of the pluripotency maintenance network in hPSCs (Varum et al., 2009; Folmes et al., 2011; Tsogtbaatar et al., 2020), and changes in glucose metabolism are among the first changes observable during differentiation (Moussaieff et al., 2015). Furthermore, differentiation to specific cell types often requires specific metabolic changes, as was observed in mesodermal (Lu et al., 2019) and neural differentiation (Lees et al., 2018). One of the ways glycolytic metabolism influences pluripotency is via sourcing of Ac-CoA for histone acetylation (Moussaieff et al., 2015), a chromatin-opening epigenetic modification regarded as one of the hallmarks of pluripotency (Mu et al., 2015; Qiao et al., 2015; Huang et al., 2016; Trisciuoglio et al., 2018). Glycolysis produces Ac-CoA by converting its end-product pyruvate to Ac-CoA through a series of reactions facilitated by PDC. Moreover, PDC is a crucial gate for glycolytic flux entrance to the TCA cycle and, subsequently, OxPhos. Considering the importance of FGF2 in pluripotency maintenance, we explored how PDH (the regulatory subunit of PDC) is controlled in response to FGF2, a cytokine commonly used in hPSCs culture, and low oxygen tensions (5% O_2_), both being known to regulate glycolytic metabolism (Papandreou et al., 2006; Fumarola et al., 2017; Liu et al., 2018).

### FGF2 promotes the use of the TCA cycle under low oxygen in hESCs to supply Ac-CoA for histone acetylation

We observed an increase in inactive (phosphorylated) PDH in 5% O_2_ compared to the 21% O_2_, suggesting reduced use of the TCA cycle and oxidative phosphorylation in hPSCs cultivated under 5% O_2_ correlating with the canonical cellular adaptation to low oxygen (Papandreou et al., 2006). Interestingly, it was FGF2 that activated PDH under 5% O_2_. This was accompanied by an increased mitochondrial membrane potential, oxygen consumption, and citrate levels, implying that FGF2 drives hPSCs to partial utilization of ethe TCA cycle and OxPhos at low oxygen levels. This is intriguing as hPSCs are usually described to prefer plain glycolysis (Folmes et al., 2011; Varum et al., 2011; Kim et al., 2015; Yu et al., 2019), and low oxygen tensions also lead to stabilization of hypoxia-inducible factors and subsequent reinforcement of glycolysis to compensate for downregulation of OxPhos due to unavailability of oxygen (Papandreou et al., 2006; Semenza, 2011). Furthermore, FGF2 signaling has been described to upregulate glycolysis in other cell types (Ward and Thompson, 2012; Yang et al., 2012; Li et al., 2016; Fumarola et al., 2017). Our data further show that active PDH is indispensable for the acetylation of histones and NANOG transcription in hPSCs, which agrees with previous findings in different cell types (Sutendra et al., 2014). Additionally, we observed that a proportion of citrate does not continue the TCA cycle in FGF2-treated cells at 5% O_2_, suggesting its utilization elsewhere, likely in histone acetylation in which Acetyl-CoA plays a central role (Moussaieff et al., 2015; Jo et al., 2020). Therefore, we propose that the increased oxygen consumption and mitochondrial membrane potential are byproducts of the PDH-mediated generation of Ac-CoA, designated for histone acetylation to support pluripotency (Moussaieff et al., 2015).

As expected, FGF2 treatment increased levels of NANOG in concordance with its pluripotency-maintaining function (Xu et al., 2005; Haghighi et al., 2018), which was further amplified in 5% O_2_ in concert with previous findings (Forristal et al., 2010). However, we did not observe a significant effect of FGF2 on PDH regulation and histone acetylation in 21% O_2_. This suggests these mechanisms are specific for the low oxygen environment, which agrees with a previous study showing that 5% O_2_ upregulates glycolysis and supports chromatin-opening epigenetic modifications in hPSCs (Lees et al., 2019). Hypoxia-inducible factors likely contrubute to this, as they mediate the switch to the glycolytic metabolism (Papandreou et al., 2006; Semenza, 2011), and H3K9Ac is dependent on HIF-1 in hPSCs (Cui et al., 2020).

### ROS directly oxidizes PDP1, elevating PDH phosphorylation

Interestingly, we observed decreased PDH phosphorylation levels after FGF2 treatment in 5% O_2_ despite detecting a higher level of its kinase PDHK1 and an unchanged amount of its phosphatase PDP1. These findings suggest that these enzyme activities are not regulated on the transcriptional level, as it occurs, for example, by TGF-β1 in fibroblasts (Smith and Hewitson, 2020). These enzymes might instead be controlled on the level of their activity, and this regulation is dependent on oxygen concentration. We have previously described how 21% O_2_ increases ROS levels enough to alter FGF2 signaling in hPSCs through oxidative inhibition of protein phosphatases (PPs) (Sommer et al., 2002; Wright et al., 2009; Fojtík et al., 2021). Here we observed a similar effect of ROS on the regulation of PDH phosphorylation. ROS have been shown to modulate the glycolytic metabolism (Liemburg-Apers et al., 2015; Molavian et al., 2016), and ROS-induced PDH phosphorylation has also been observed (Cesi et al., 2017). Here we show, in concert with these data, that ROS inactivate PDP1 in hPSCs in low O_2_ by direct oxidation, leading to a quantitative decrease in PDH phosphorylation and subsequent downregulating of histone acetylation and NANOG expression. These observations correspond with a recent study describing that the PDP1 activation of PDH is required for the acetylation of histones in cancer cell lines in hypoxia (Karagiota et al., 2022). A limitation of our study is the lack of data on the activity-enhancing tyrosine phosphorylation of PDHK1 (Hitosugi et al., 2011). However, ROS-mediated activation of PDHK1 has been previously observed (Cesi et al., 2017), and it was possibly also upregulated because of the decreased phosphatase activity (Sommer et al., 2002; Wright et al., 2009), which would work in synergy with the inactivation of PDP1 observed by us.

PDPs belong to a Ser/Thr protein phosphatase 2C family, and Ser/Thr PPs are usually described to have only low or no sensitivity to oxidation (Sommer et al., 2002). However, some researchers previously observed their oxidation (Pieri et al., 2003) and even argued that failures to make this observation were due to experiments performed at high O_2_ levels (Wright et al., 2009), which were shown to inhibit Ser/Thr PPs (Nyunoya et al., 2005). Our results add to the body of literature describing Ser/Thr PPs’ sensitivity to oxidation and support the concept that 21% O_2_ induces ROS levels high enough to inactivate Ser/Thr PPs, thus obscuring the effects of exogenous oxidants.

There are three phosphorylation sites on PDH, and phosphorylation of either of them results in PDH inactivation. The Ser^293^ is the most rapidly phosphorylated site (Yeaman et al., 1978) and the most abundantly phosphorylated site across tissues (Rardin et al., 2009). Four isoforms of PDHK (1-4) are described to date, having distinct tissue and PDH phosphorylation site-specificity, but PDHK1 is the only isoform capable of phosphorylating all three PDH sites with the highest affinity for Ser^293^ (Yeaman et al., 1978). Moreover, PDHK1 upregulation has been described in hPSCs (Varum et al., 2011). Therefore, we focused on PDH Ser^293^ phosphorylation and PDHK1 when studying the regulation of PDH.

To induce ROS in this study, we used 5μM H_2_O_2_, the upper limit of H_2_O_2_ detected in plasma (Forman et al., 2016). After perturbing the cell (Lyublinskaya and Antunes, 2019), H_2_O_2_ of this concentration should increase its intracellular levels slightly above the detected physiological levels (Sies, 2017) or even 5- to 7-fold according to a newer study (Lyublinskaya and Antunes, 2019). Therefore, the molecular changes we observe are direct effects, not oxidative stress artifacts. This is also supported by the fact that the increase in ROS caused by 21% O_2_ seems to be sufficient to modulate the activity of PDP1 and PI3K/AKT signaling (Fojtík et al., 2021).

Furthermore, FGF2 treatment or deprivation for 24 hours did not affect mitochondrial morphology, as mitochondria remained small, perinuclear, and with undeveloped cristae, typical for hPSCs (Folmes et al., 2011; Varum et al., 2011). FGF2 signaling also did not affect mitochondrial morphology during their enforced maturation induced by treatment with PDHKs inhibitor dichloroacetate, which also testified to the metabolic plasticity of our hPSCs. These observations suggest that the 24h withdrawal of FGF2 that was used in our experiments does not change hPSCs bioenergetics towards the differentiated type (Folmes et al., 2011), and we are not making comparisons with cells of different metabolic phenotypes.

### MEK1/2-ERK1/2 participates in PDH activity regulation by FGF2 by downregulating ROS

To dissect how FGF2 signaling regulates PDH phosphorylation, we looked at its two main downstream pathways involved in regulating energy metabolism – MEK1/2-ERK1/2 and PI3K/AKT. MEK1/2-ERK1/2 signaling was shown to upregulate glycolysis and promote the Warburg effect by phosphorylation and nuclear translocation of PKM2 (Yang et al., 2012). Contrary to this, our results suggest that MEK1/2-ERK1/2 signaling promotes attenuation of ROS (Cesi et al., 2017; Fojtík et al., 2021), alleviating PDP1 inhibition and ultimately leading to dephosphorylation of PDH in 5% O_2_. At the same time, we show that MEK1/2-ERK1/2 increases mitochondrial membrane potential, suggesting utilization of oxidative phosphorylation, which agrees with elevated respiration in FGF2-treated cells. The difference between our and previous observations might be caused by using different cell lines at different oxygen concentrations, as we saw this phenotype exclusively in 5% O_2_.

ERK1/2 could downregulate ROS by phosphorylation of HIFs, which increases their transcriptional activity (Richard et al., 1999; Mylonis et al., 2008), consequently increasing the production of GSH and NADPH (Samanta and Semenza, 2017), core elements of cellular antioxidant system. This is supported by our previous observation that in hPSCs, MEK1/2 inhibition in 21% O_2_ does not affect ROS levels (Fojtík et al., 2021).

We have previously described that increased ROS levels upregulate PI3K/AKT (Fojtík et al., 2021), which could then lead to the phosphorylation of PDH (Cerniglia et al., 2015) as PI3K/AKT pathway upregulates glycolysis through its downstream effectors FoxO and mTOR (Xie et al., 2019). However, our experiments with the AKT activator show that PI3K/AKT does not increase PDH phosphorylation in hPSCs at 5% O_2_.

Although our data show that the MEK1/2-ERK1/2 pathway controls PDH phosphorylation via the regulation of ROS levels, it seems to regulate histone acetylation also by other mechanisms different from the ROS-mediated phosphorylation of PDH. MEK1/2 inhibition downregulated AcH3 stronger than the plain absence of FGF2. Such mechanisms have been widely studied in the context of memory formation (Levenson et al., 2004). One of the proposed modes of action is the direct ERK1/2-mediated phosphorylation of histones the and subsequent recruitment of the Gcn5 histone acetyltransferase (Cheung et al., 2000).

Effective and safe use of hPSCs in basic research, disease modeling, and potentially cell replacement therapies require precise control over their fate as hPSCs have been shown to fluctuate between different degrees of pluripotent and primed states (Kalmar et al., 2009; Nguyen et al., 2018; Hayashi et al., 2019). This study complements our previous work (Fojtík et al., 2021) showing that control over ROS homeostasis is also relevant for pluripotency maintenance, and we expand it to the regulation of PDH, a gatekeeper of Ac-CoA production, which is required for maintenance of pluripotent epigenetic landscape (Jo et al., 2020). Furthermore, we describe a novel role of FGF2-MEK1/2-ERK1/2 signaling in regulating PDP1 activity through ROS attenuation, resulting in increased NANOG transcription. Our results suggest that implementing control over PDH activity and ROS levels might greatly enhance current cultivation or even differentiation protocols, as ROS are implicated in supporting cardiac and neuronal differentiation (Bell et al., 2015; Momtahan et al., 2019). Furthermore, our insights into the ROS-mediated regulation of PDP1 might also find use in cancer biology as MEK1/2-ERK1/2 targeting treatment (Cesi et al., 2017) and potentially other types of treatment that lead to increased levels of ROS can potentially steer the energy metabolism towards the unfavorable aerobic glycolysis by inhibiting PDH.

## Experimental procedures

A complete list of experimental procedures can be found in Supplemental Figure 1.

## Resource availability

### Corresponding author

Further information and requests for resources and reagents should be directed to and will be fulfilled by the corresponding author, Vladimir Rotrekl

### Materials availability

This study generated no new unique cell lines or reagents.

### Data and code availability

Datasets used to generate figures in this publication are available at DOI: 10.6084/m9.figshare.21883026. For other source data please contact corresponding author Vladimir Rotrekl.

## Statistical analysis

The number of independent experiments is indicated in the figure legends (N). Arithmetical means, standard error of the mean (SEM), and statistical tests were calculated using the GraphPad Prism 8 software (GraphPad Software, La Jolla, CA, USA). Statistical significances were determined using a one sample t-test (theoretical mean = 1) when making a comparison with a control sample (value = 1) and a paired two-tailed t-test when comparing other samples. The use of different tests or variation measurements (Figures 2A and 4B) are denoted in corresponding figure legends. Values used to construct the graphs can be found among shared datasets (DOI: 10.6084/m9.figshare.21883026). Asterisks indicate a significant difference: *p<0.05; **p<0.01; ***p<0.001.

## Acknowledgment and funding

This work was supported by funds from the Faculty of Medicine, Masaryk University, to a junior researcher Jana Gregorová (ROZV/28/LF10/2020), by the Specific University Research Grant (MUNI/A/1275/2022), by the European Regional Development Fund - Project ENOCH (No. CZ.02.1.01/0.0/0.0/16_019/0000868) and project National Institute for Research of Metabolic and Cardiovascular Diseases (Programme EXCELES, ID Project No. LX22NPO5104) – Funded by the European Union – Next Generation EU. This work was also supported by project nr. LX22NPO5107 (MEYS): Financed by EU – Next Generation EU and has received funding from the European Union’s Horizon 2020 research and innovation programme under grant agreement No 857560. This publication reflects only the author’s view and the European Commission is not responsible for any use that may be made of the information it contains.

We want to thank Jana Okunkova for her help with cell culture and Deborah Beckerova for her help with RNA isolation.

## Author contributions

Conceptualization, S.U., V.R., and P.F.; Methodology, P.F., V.R., J.G., and O.P.; Formal analysis, P.F., and A.S.; Investigation, P.F., M. Sen., K.H., J.R., M. Ste., M. Sed., A.S., J.G., O.S.; Resources, V.R., P.S., M.Sed., A.H., D.B., J.G.; Writing – original draft, P.F., V.R.; Funding acquisition, V.R., J.G., D.B.

