## Supplemental information for "FGF2-induced Redox Signaling: A Mechanism Regulating Pyruvate Dehydrogenase Driven Histone Acetylation and NANOG Upregulation"

### Supplemental Figures and Tables

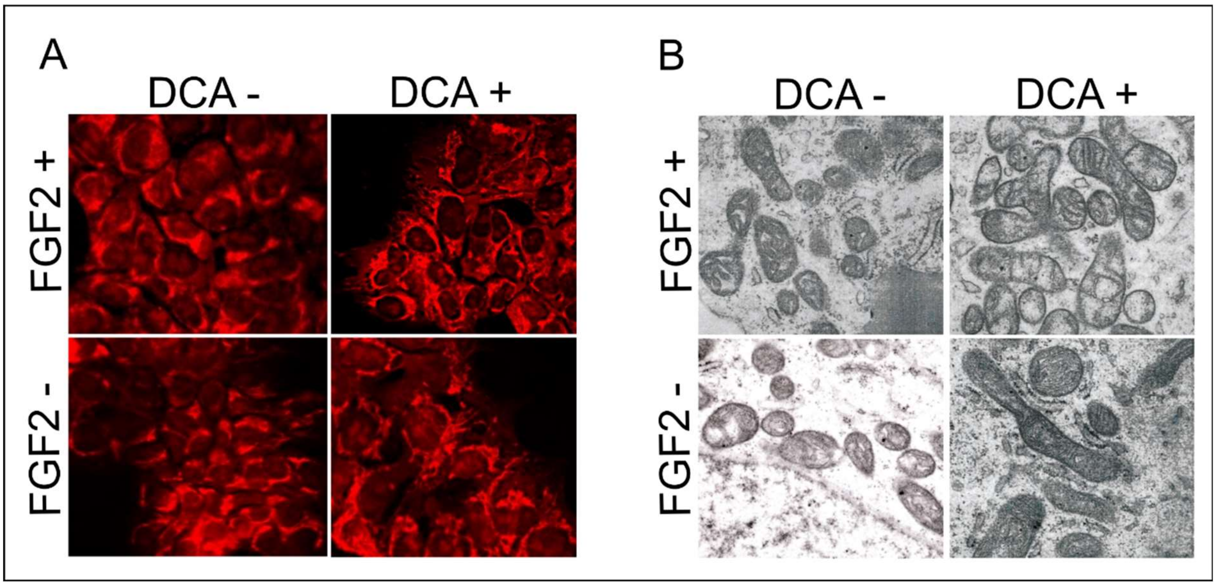

**Figure S1**. **24-hour FGF2 starvation does not affect mitochondrial structure and development.**

**A)** MitotrackerRed CMXRos staining of mitochondria in CCTL14 hPSCs.

**B)** Transmission electron microscopy of CCTL14 hPSCs with detail on the mitochondria.

**Table S1. List of compounds used to treat the hPSCs in this study.** This table is related to the Supplementary experimental procedures – Cell culture.

| Name | Abbreviation | Final concentration | Effect | Cat. # | Manufacturer |
|---|---|---|---|---|---|
| PD0325901 | PD03 | 0.2μM | MEK1/2 inhibition | PZ0162 | Sigma-Aldrich |
| SC79 | SC79 | 10μM | AKT activation | S7863 | Selleck Chemicals |
| CPI-613 | CPI | 10μM | PDH inhibition | S2776 | Selleck Chemicals |
| Hydrogen peroxide | $H_2O_2$ | 5mM | ROS induction | H1009 | Sigma-Aldrich |
| Glutathione (reduced) | GSH | 5mM | ROS quenching | G6013 | Sigma-Aldrich |
| Sodium citrate | Cit | 5mM | Citrate supplementation | W302600 | Sigma-Aldrich |

**Table S2. List of primary and secondary antibodies used in Western blotting.** This table is related to the Supplementary experimental procedures – Western blotting.

| **Primary antibodies** | | | | |
|---|---|---|---|---|
| **Target** | **Raised in** | **Dilution** | **Cat. #** | **Manufacturer** |
| ERK1/2 | Rabbit | 1:1000 | 9102 | Cell Signaling Technology |
| pERK1/2 | Rabbit | 1:1000 | 9101 | Cell Signaling Technology |
| AKT | Rabbit | 1:1000 | 9272 | Cell Signaling Technology |
| pAKT | Rabbit | 1:1000 | 9271 | Cell Signaling Technology |
| α-Tubulin | Mouse | 1:2000 | 11-250-C100 | Exbio |
| PCNA | Rabbit | 1:2000 | HPA030522 | Sigma-Aldrich |
| AcH3 | Rabbit | 1:2000 | 47915 | Abcam |
| PDH | Rabbit | 1:1000 | 2784 | Cell Signaling Technology |
| pPDH ($Ser^{293}$) | Rabbit | 1:500 | AP1062 | Calbiochem |
| PDHK1 | Rabbit | 1:1000 | 3820 | Cell Signaling Technology |
| PDP1 | Rabbit | 1:1000 | 65575 | Cell Signaling Technology |
| Vinculin | Rabbit | 1:2000 | 13901 | Cell Signaling Technology |
| Nanog | Rabbit | 1:500 | Sc-33759 | Santa Cruz Biotechnology |
| **Secondary antibodies** | | | | |
| **Name** | **Raised in** | **Dilution** | **Cat. #** | **Manufacturer** |
| anti-rabbit IgG-HRP | Goat | 1:3000 | 7074 | Cell Signaling Technology |
| anti-mouse IgG-HRP | Goat | 1:5000 | 12-349 | Merck Millipore |

**Table S3. List of primers used in qRT-PCR.** This table is related to the Supplementary experimental procedures – RNA isolation and quantitative real-time PCR (qRT-PCR).

| | | |
|---|---|---|
| GAPDH | forward | 5'-AGCCACATCGCTCAGACACC-3' |
| | reverse | 5'-GTACTCAGCGCCAGCATCG-3' |
| Nanog | forward | 5'-CCTATGCCTGTGATTTGTGG-3' |
| | reverse | 5'-CTGGGACCTTGTCTTCCTTT-3' |

**Table S4. SRM transitions (Q1 → Q3, collision energy, and dwell time) of compounds analyzed in the intracellular content.** This table is related to the Supplementary experimental procedures – LC-MS.

| Compound | Polarity | Q1 (m/z) | Q3 (m/z) | CE (eV) | Dwell time (ms) |
|---|---|---|---|---|---|
| 2-OG | – | 145.0 | 101.1 | 40 | 10.0 |
| | – | 145.0 | 57.3 | 7 | 10.0 |
| 3-Phosphoglycerate | – | 185.0 | 97.1 | 9 | 7.5 |
| | – | 185.0 | 79.1 | 24 | 7.5 |
| | – | 185.0 | 167.0 | 3 | 7.5 |
| 6-Phosphogluconate | – | 275.1 | 97.1 | 10 | 12.5 |
| | – | 275.1 | 79.1 | 26 | 12.5 |

| | | | | | |
|---|---|---|---|---|---|
| | – | 275.1 | 177.1 | 10 | 12.5 |
| | – | 275.1 | 99.1 | 18 | 12.5 |
| 8-hydroxy deoxyguanosin | + | 284.0 | 168.1 | -11 | 11.3 |
| | + | 284.0 | 117.2 | -12 | 11.3 |
| Acetyl-CoA | + | 810.0 | 302.8 | -26 | 5.0 |
| | + | 810.0 | 427.9 | -24 | 5.0 |
| | + | 810.0 | 201.0 | -29 | 5.0 |
| | + | 810.0 | 135.9 | -47 | 5.0 |
| Adenosine | + | 268.1 | 136.1 | -13 | 10.0 |
| | + | 268.1 | 119.1 | -44 | 10.0 |
| ADP | + | 428.0 | 136.1 | -15 | 20.0 |
| Ala | + | 90.1 | 44.1 | -9 | 11.3 |
| | + | 90.1 | 43.2 | -9 | 11.3 |
| AMP | – | 346.0 | 210.9 | 10 | 16.7 |
| | – | 346.0 | 79.2 | 25 | 16.7 |
| | – | 346.0 | 134.1 | 23 | 16.7 |
| Arg | + | 175.2 | 70.1 | -25 | 11.3 |
| | + | 175.2 | 60.1 | -25 | 11.3 |
| Ascorbic acid | – | 175.0 | 87.2 | 14 | 6.7 |
| | – | 175.0 | 115.1 | 10 | 6.7 |
| | – | 175.0 | 71.2 | 7 | 6.7 |
| Asn | + | 133.1 | 74.2 | -12 | 7.5 |
| | + | 133.1 | 87.2 | -7 | 7.5 |
| | + | 133.1 | 116.1 | -7 | 7.5 |
| Asp | – | 132.0 | 88.2 | 7 | 11.3 |
| | – | 132.0 | 115.1 | 7 | 11.3 |
| ATP | – | 506.0 | 159.0 | 20 | 16.7 |
| | – | 506.0 | 115.0 | 20 | 16.7 |

| | | | | | |
|---|---|---|---|---|---|
| | – | 506.0 | 408.0 | 20 | 16.7 |
| | + | 508.0 | 136.1 | -20 | 22.5 |
| B12 | + | 678.6 | 359.0 | -25 | 11.3 |
| | + | 678.6 | 456.6 | -30 | 11.3 |
| cAMP | – | 328.1 | 134.1 | 20 | 16.7 |
| | – | 328.1 | 79.2 | 26 | 16.7 |
| | – | 328.1 | 107.2 | 40 | 16.7 |
| | + | 330.2 | 136.1 | -22 | 16.7 |
| | + | 330.2 | 119.1 | -44 | 16.7 |
| | + | 330.2 | 312.0 | -12 | 16.7 |
| Citric acid | – | 191.0 | 111.1 | 9 | 7.5 |
| | – | 191.0 | 87.2 | 17 | 7.5 |
| | – | 191.0 | 85.2 | 11 | 7.5 |
| Citrulline | + | 176.1 | 159.1 | -6 | 6.7 |
| | + | 176.1 | 70.3 | -18 | 6.7 |
| | + | 176.1 | 113.2 | -12 | 6.7 |
| CMP | + | 324.0 | 112.1 | -10 | 7.5 |
| | + | 324.0 | 279.1 | -10 | 7.5 |
| | + | 324.0 | 307.1 | -5 | 7.5 |
| | – | 322.0 | 210.9 | 10 | 7.5 |
| | – | 322.0 | 79.1 | 24 | 7.5 |
| | – | 322.0 | 97.1 | 21 | 7.5 |
| Creatinine | + | 114.1 | 44.4 | -15 | 10.0 |
| | + | 114.1 | 86.2 | -8 | 10.0 |
| Cys | + | 122.0 | 59.3 | -19 | 7.5 |
| | + | 122.0 | 76.2 | -10 | 7.5 |
| | + | 122.0 | 105.1 | -7 | 7.5 |
| Folic acid | + | 442.1 | 295.0 | -22 | 16.7 |

| | | | | | |
|---|---|---|---|---|---|
| | + | 442.1 | 176.0 | -38 | 16.7 |
| | + | 442.1 | 120.1 | -32 | 16.7 |
| Fru-1,6-BisP | – | 339.0 | 97.1 | 14 | 16.7 |
| | – | 339.0 | 79.1 | 29 | 16.7 |
| | – | 339.0 | 240.9 | 7 | 16.7 |
| Fumarate | – | 115.0 | 71.2 | 4 | 20.0 |
| Glu | + | 148.1 | 84.2 | -13 | 7.5 |
| | + | 148.1 | 130.1 | -6 | 7.5 |
| | + | 148.1 | 102.2 | -9 | 7.5 |
| Glucuronic acid | – | 193.0 | 113.1 | 6 | 16.7 |
| | – | 193.0 | 59.3 | 13 | 16.7 |
| | – | 193.0 | 73.2 | 10 | 16.7 |
| Gly | + | 76.0 | 30.5 | -7 | 10.0 |
| | + | 76.0 | 48.4 | -4 | 10.0 |
| GMP | + | 364.0 | 152.1 | -11 | 6.7 |
| | + | 364.0 | 135.0 | -35 | 6.7 |
| | + | 364.0 | 110.2 | -42 | 6.7 |
| GSH | + | 308.1 | 76.2 | -15 | 7.5 |
| | + | 308.1 | 84.2 | -15 | 7.5 |
| | + | 308.1 | 161.9 | -15 | 7.5 |
| Guanosine | + | 284.1 | 152.1 | -10 | 7.5 |
| | + | 284.1 | 135.1 | -36 | 7.5 |
| | + | 284.1 | 110.1 | -37 | 7.5 |
| Hex-6-P | – | 259.0 | 97.1 | 10 | 11.3 |
| | – | 259.0 | 79.1 | 29 | 11.3 |
| Hexose | – | 179.1 | 59.3 | 15 | 7.5 |
| | – | 179.1 | 89.2 | 4 | 7.5 |
| | – | 179.1 | 119.1 | 4 | 7.5 |

| | | | | | |
|---|---|---|---|---|---|
| His | + | 156.2 | 110.0 | -13 | 11.3 |
| | + | 156.2 | 83.1 | -13 | 11.3 |
| Homocysteine | + | 136.0 | 90.2 | -11 | 10.0 |
| | + | 136.0 | 118.1 | -3 | 10.0 |
| Ile | + | 132.1 | 86.1 | -15 | 11.3 |
| | + | 132.1 | 69.1 | -15 | 11.3 |
| Lactate | – | 89.0 | 43.4 | 6 | 10.0 |
| | – | 89.0 | 45.3 | 9 | 10.0 |
| Leu | + | 132.1 | 86.1 | -15 | 11.3 |
| | + | 132.1 | 69.1 | -15 | 11.3 |
| Lys | + | 147.2 | 84.1 | -17 | 22.5 |
| Malate | – | 133.0 | 115.0 | 4 | 6.7 |
| | – | 133.0 | 71.2 | 13 | 6.7 |
| | – | 133.0 | 73.2 | 16 | 6.7 |
| Met | + | 150.0 | 56.3 | -14 | 11.3 |
| | + | 150.0 | 104.2 | -9 | 11.3 |
| Methylcobalamine | + | 672.8 | 665.0 | -5 | 6.7 |
| | + | 672.8 | 147.1 | -43 | 6.7 |
| | + | 672.8 | 359.0 | -26 | 6.7 |
| Methyl-tetrahydrofolate | + | 460.2 | 313.1 | -18 | 16.7 |
| | + | 460.2 | 180.0 | -36 | 16.7 |
| | + | 460.2 | 194.1 | -32 | 16.7 |
| NAD+ | + | 664.1 | 428.0 | -25 | 11.3 |
| | + | 664.1 | 348.1 | -35 | 11.3 |
| NADH | + | 666.1 | 514.1 | -25 | 7.5 |
| | + | 666.1 | 428.0 | -15 | 7.5 |
| | + | 666.1 | 302.0 | -20 | 7.5 |
| NADP+ | + | 744.1 | 136.2 | -50 | 12.5 |

| | | | | | |
|---|---|---|---|---|---|
| | + | 744.1 | 507.9 | -26 | 12.5 |
| | + | 744.1 | 604.1 | -16 | 12.5 |
| | + | 744.1 | 622.1 | -8 | 12.5 |
| | + | 372.0 | 123.1 | -8 | 11.3 |
| | + | 372.0 | 136.1 | -21 | 11.3 |
| NADPH | – | 744.1 | 159.0 | 47 | 12.5 |
| | – | 744.1 | 79.1 | 53 | 12.5 |
| | – | 744.1 | 621.9 | 8 | 12.5 |
| | – | 744.1 | 396.8 | 32 | 12.5 |
| | – | 371.5 | 79.1 | 24 | 12.5 |
| | – | 371.5 | 134.1 | 13 | 12.5 |
| | – | 371.5 | 304.0 | 5 | 12.5 |
| | – | 371.5 | 158.9 | 20 | 12.5 |
| OH-Pro | + | 132.0 | 86.2 | -10 | 7.5 |
| | + | 132.0 | 68.3 | -16 | 7.5 |
| | + | 132.0 | 41.4 | -29 | 7.5 |
| Orn | + | 133.2 | 70.1 | -17 | 11.3 |
| | + | 133.2 | 116.0 | -17 | 11.3 |
| Panthotenic acid | + | 220.1 | 98.0 | -25 | 11.3 |
| | + | 220.1 | 124.1 | -22 | 11.3 |
| PEP | – | 167.0 | 79.1 | 6 | 20.0 |
| Phe | + | 166.2 | 120.0 | -13 | 11.3 |
| | + | 166.2 | 77.1 | -13 | 11.3 |
| Pro | + | 116.1 | 70.1 | -17 | 11.3 |
| | + | 116.1 | 43.2 | -17 | 11.3 |
| Pyridoxine | + | 170.1 | 134.1 | -25 | 11.3 |
| | + | 170.1 | 106.1 | -26 | 11.3 |
| Ribose-5-P | – | 229.0 | 97.1 | 8 | 11.3 |

| | | | | | |
|---|---|---|---|---|---|
| | – | 229.0 | 79.1 | 24 | 11.3 |
| SAH | + | 385.1 | 136.1 | -19 | 11.3 |
| | + | 385.1 | 134.1 | -19 | 11.3 |
| SAM | + | 399.1 | 250.1 | -12 | 7.5 |
| | + | 399.1 | 136.1 | -19 | 7.5 |
| | + | 399.1 | 298.1 | -11 | 7.5 |
| Ser | + | 106.0 | 60.3 | -9 | 7.5 |
| | + | 106.0 | 42.4 | -19 | 7.5 |
| | + | 106.0 | 88.2 | -6 | 7.5 |
| Succinate | – | 117.0 | 73.2 | 8 | 10.0 |
| | – | 117.0 | 99.1 | 6 | 10.0 |
| Sulphate | – | 97.1 | 80.1 | 21 | 20.0 |
| Taurine | – | 124.0 | 80.1 | 17 | 20.0 |
| Tetrahydrofolate | + | 446.2 | 166.1 | -11 | 16.7 |
| | + | 446.2 | 299.1 | -16 | 16.7 |
| | + | 446.2 | 178.1 | -11 | 16.7 |
| Thr | + | 120.1 | 74.3 | -8 | 11.3 |
| | + | 120.1 | 56.3 | -13 | 11.3 |
| Trp | + | 205.1 | 188.0 | -5 | 11.3 |
| | + | 205.1 | 146.0 | -5 | 11.3 |
| Tyr | + | 182.2 | 136.0 | -9 | 11.3 |
| | + | 182.2 | 165.0 | -9 | 11.3 |
| Uridine | + | 245.1 | 113.1 | -6 | 11.3 |
| | + | 245.1 | 70.2 | -24 | 11.3 |
| Val | + | 118.1 | 72.3 | -8 | 11.3 |
| | + | 118.1 | 55.4 | -17 | 11.3 |

### Supplemental Experimental Procedures

#### Cell culture

Experiments were performed using hESCs line CCTL12 and CCTL14 and the human iPSCs line AM13, previously characterized previously by Adewumi and Krutá et. al.[1,2]. Long-term cultivation of hPSCs was performed on mitotically inactivated mouse embryonic fibroblast (MEF) from the CD1 and CF1 mouse strain in the human embryonic stem cell media (hES) containing the Dulbecco's modified Eagle medium (DMEM)/F12 (Thermofisher Scientific, Waltham, MA, USA, 21331-020) supplemented with 15% (vol/vol) knockout serum replacement (Thermofisher Scientific, 10828-028), nonessential amino acids (Thermofisher Scientific, 11140-035), 4 ng/ml FGF-2 (PeproTech, Cranbury, NJ, USA, 100-18B), L-glutamine, 0.5% (vol/vol; Biosera, Boussens, France, XC-T1715), penicillin-streptomycin (Biosera, XC-A4122) and 2-mercaptoethanol (Sigma-Aldrich, St. Louis, MO, USA, M3148) in a colony type culture. For experimental procedures, hPSCs were cultured feeder-free on Matrigel hESC-qualified Matrix (Matrigel; Corning, NY, USA, 354277) coated dishes and cultivated in MEF-conditioned hES media (CM+) supplemented with FGF2 (10 ng/ml) in a monolayer-type culture. hES media was conditioned on mitotically inactivated MEF for 24 hours, and the same dish containing MEF was used seven times, then all seven batches were mixed, supplemented with L-Glutamine (0.5% vol/vol) and FGF2 (10 ng/ml) and filtered. MEF-conditioned media without FGF2 (CM-) was prepared using hES media missing FGF2, and the CM- was not further supplemented with FGF2. hPSCs maintained on matrigel-coated dishes were cultivated for a maximum of seven passages. For cultivation in 5% $O_2$, cells were kept in the MCO-18M multigas incubator (Sanyo, Moriguchi, Osaka, Japan). On a feeder culture, hPSCs were passaged mechanically. On a matrigel culture, hPSCs were passaged by dissociation using TrypLE Express Enzyme (TrypLE; Thermofisher scientific, 12605010), spun 200g/4min/4°C, and resuspended in a fresh media. Experimental treatments are listed in Table S1.

#### Gene expression silencing

Endoribonuclease-prepared small interfering RNAs (esiRNA) against PDP1 (10nmol; Sigma-Aldrich, NM_018444: SASI_Hs01_00128068) were introduced into the cells using Lipofectamine 2000 Transfection Reagent (Thermofisher Scientific, 11668-037) according to the manufacturer's manual. Successful downregulation (approximately 48 hours after transfection and 2 hours after media change) was confirmed using Western blot (WB).

#### Western blotting

hPSCs cultivated on matrigel coated dishes to the maximum of 70% confluence were washed three times with 1x phosphate-buffered saline (PBS) and lysed on ice with 1% SDS lysis buffer (50mM Tris-HCl, 1% SDS, pH 6,8). Protein concentrations were quantified using the DC Protein Assay (Bio-Rad, Hercules, CA, USA, 5000111) measured in triplicates on DTX 880 Multimode Detector (Beckman Coulter, Brea, CA, USA). Protein concentrations were then adjusted to 1 mg/ml, 10x Laemmli buffer was added, and the lysates were briefly boiled. SDS-PAGE was performed using either 8% or 10% polyacrylamide gels, 15 µg of total protein was loaded, and gel electrophoresis ran at 140V/70 min. Proteins were then transferred to Immobilon-P PVDF membrane (Merck Millipore, Burlington, MA, USA, IPVH00010) 100V/60 min.

The membranes were blocked in 5% dried milk in tris buffered saline containing 0,1% Tween-20 (TBS-T) for 1 hour and incubated overnight with antibodies diluted in the respective blocking solutions at 4°C. The following day the membranes were washed 3x 15 min with TBS-T and incubated with secondary antibodies diluted in the respective blocking solution for 1 hour at room temperature, followed by 5x 10 min washes with TBS-T. Immobilon Western Chemiluminescent HRP Substrate (Merck Millipore, P90720) was used as a substrate for the luminescence reaction. Image developing was performed using the G:Box Chemi device (SYNGENE, Bangalore, India). The obtained images were adjusted using the GIMP2 software and analyzed in ImageJ.

For the detection of PDP1 oxidation-induced changes, cells were harvested in native lysis buffer (100mM Tris pH7, 150mM NaCl, 1mM EDTA, 0,1% Triton X-100) supplemented with the cOmplete Mini Protease Inhibitor Cocktail (Roche, Basel, Switzerland; 11836153001) and 50mM N-Ethylmaleimide (Sigma-Aldrich; 04259). Samples were sonicated, mixed with a non-reducing loading buffer, and resolved using 8% SDS-PAGE at 4°C. Protein transfer and detection were performed as described above.

Primary and secondary antibodies used in this study are listed in Table S2. WB quantification analysis was performed using the ImageJ software as described before[3]. Proteins of interest were normalized to loading control.

**Analysis of reactive oxygen species (ROS) generation**
Cells grown on Matrigel-coated coverslips were deprived of FGF2 24 hours before treatment. Fifty minutes before the end of the treatment CellROX Green (5µM; ThermoFisher Scientific, C10444) was added. Afterwards, the cells were washed 3x in PBS on ice and fixed with 4% paraformaldehyde (Sigma-Aldrich, 158127) for 30 minutes. Snapshots of the approximately same-sized cell clusters were taken with constant exposure time using the fluorescent LSM700 microscope (×40 1.3 oil differential interference contrast lens; Carl Zeiss, Oberkochen, Germany) within 6 hours of fixation. ROS levels were determined as fluorescence divided by the fluorescence area in raw images using the ImageJ software.

**RNA isolation and quantitative real-time PCR (qRT-PCR)**
Total RNA was isolated using the RNA Blue reagent (Top-Bio, Czech Republic) according to the manufacturer's protocol. mRNA concentration and purity were determined using the NanoDrop device (NanoDrop Technologies, Wilmington, Germany). 2 µg of total RNA were transcribed into cDNA using the Moloney Mouse Leukemia Virus (M-MLV) reverse transcriptase (Invitrogen, Carlsbad, CA, USA) and Oligo(dT) primers (Thermo Fisher Scientific Inc., USA) at 37 °C for 1 h followed by 5 min at 85 °C. qRT-PCR was performed using the LightCycler®480 DNA SYBR Green I Master (Roche) in a Light Cycler 480 instrument. The obtained data were normalized to Glyceraldehyde 3-phosphate dehydrogenase (GAPDH) mRNA expression and are presented as $2^{-\Delta cq}$. Primer sequences are listed in Table S3.

**Mitochondrial membrane potential measurement**
The mitochondrial membrane potential was visualized using the tetramethylrhodamine methyl ester (TMRM; Invitrogen, T668) fluorescent probe. TMRM is a cationic, membrane-penetrable molecule that enters the mitochondrial intermembrane space based on the proton gradient, and therefore, its fluorescence is proportional to the proton potential on the membrane. It has an excitation maximum in the green part and an emission in the yellow part of the light spectrum. TMRM was added to complete media in the final 20nM concentration and incubated with the cells for 20 minutes at 37°C. Cells were then washed with 1x PBS and harvested using TrypLE. The supernatant was discarded, and the pellet was resuspended in 0.5 ml ice-cold PBS. The samples were handled on ice and immediately analyzed using the Beckman Coulter Cytomics FC 500 flow cytometer (Beckman Coulter, Brea, CA, USA). Using the forward and side scatter, the populations of singlets were selected, and in every measurement, at least 10 000 singlet events were recorded. Fluorescence data were recorded on logarithmic axes, and mean fluorescence was determined. Further data analysis was performed using the FlowJo 7.2.2. software (FlowJo, Ashland, OR, USA).

**Mitotracker Red CMXRos staining**
Mitochondria of hPSCs cultivated on Matrigel-coated coverslips were visualized using Mitotracker Red CMXRos (MTT; Life Technologies) according to the manufacturer's manual. Briefly, 1mM stock solution was diluted to 250 µM in DMEM/F12 and added to cell culture to a final concentration of 25 nM for 20 minutes. After that, hPSCs were washed 5 times with preheated DMEM/F12; dishes were transferred on ice, washed 3 times with 1x PBS, and fixed with 4% paraformaldehyde for 30 minutes on ice in the dark and mounted. Snapshots of the approximately same-sized cell clusters were taken using the fluorescent LSM700 microscope (×40 1.3 oil differential interference contrast lens; Carl Zeiss, Oberkochen, Germany)

**Electron microscopy**
CCTL14 hPSCs were seeded on matrigel-coated dishes, grown for 24 h with the corresponding treatments, and harvested using TrypLE, washed with 1× PBS, fixed, and prepared as described before [4]. Ultrathin sections were prepared on Leica EM UC6 ultramicrotome, stained with uranyl acetate and Reynold's lead citrate, and examined with the FEI Morgagni 286(D) TEM.

**Oxygen consumption measurement**
To measure the dynamics of oxygen consumption, hPSCs were grown in a monolayer on Matrigel-coated Petri dishes with 2 ml of conditioned medium. Oxygen level was followed continuously with miniaturized Clark-type sensors (BVT Technologies, Brno, Czech Republic) with polypropylene membrane mounted on an exchangeable holder. The measuring part of the sensor was embedded in the lid of the Petri dish at a 2 mm distance from the cell culture. Two individual sensors were simultaneously connected to the QuadStat EA164 electrochemical analyzer (eDAQ, Denistone East, Australia). The working potential was -650 mV vs. the Ag/AgCl electrode as a reference. The data sampling interval was 1 s using the Chart software (ver. 5.5.16) from eDAQ. Before measurements, the sensors were calibrated using sodium sulphite solutions. After a wash with water and 70% ethanol, the

sensors were placed in a fresh cultivation medium and allowed to stabilize for 30 min before transfer to the Petri dish with cell culture. All measurements were performed at atmospheric pressure (101 kPa) and 37 °C. To determine the oxygen level ($O_m$, μM), the measured current ($C_m$, mA) was normalized by the current measured in distilled water after equilibration at 37 °C ($C_w$, mA) and then multiplied by the concentration of $O_2$ (Ow, μM) in distilled water under corresponding environmental c conditions (Equation 1):

$$O_m = \frac{C_m}{C_w} O_w$$

The oxygen concentration in distilled water is equal to 212 μM (from the IUPAC tables, under water-saturated air at a total pressure of 101 kPa and temperature of 37°C). Linear regression of Om data was done, and the value of the regression line slope represents the oxygen consumption rate.

**Liquid chromatography coupled with mass spectrometry (LC-MS)**
Cells cultivated in 21% and 5% $O_2$ were treated with FGF2 (10 ng/ml) after 24 h of FGF2 deprivation and then for another 2 h. Then, they were washed with ice-cold PBS on ice (4°C), collected in LC-MS buffer (60% acetonitrile, 30% methanol, 10% $H_2O$), freeze-lysed at -80°C and analyzed within 2 hours of this process.

Metabolite standards, acetonitrile (ACN), methanol (MeOH), acetic acid, formic acid, and ammonium hydroxide were purchased from Sigma-Aldrich (Prague, Czech Republic). All reagents used were of an analytical grade except for methanol and acetonitrile, which were of an LC-MS grade. Water used for LC-MS was of an ultra-pure grade supplied by an in-house Milli-Q system (Millipore, MA, USA).

Metabolite standards were dissolved in water to prepare stock solutions (0,1 mg/ml). The stock solutions were stored at − 80 °C. Working standard solutions were made by diluting the stock solutions with ACN. The final concentration of the metabolites was 10 μg/ml. The metabolite mixture was used to obtain LC MS characteristics of each metabolite, e.g., retention time, peak shape, MS/MS behavior, or sensitivity, which could be compared to peaks obtained from LC-MS of the lysed cells. Replicates (n = 3) of cells were mixed with 1 ml of 90% ACN, shortly centrifuged, and 20 μL of the supernatant was injected onto the analytical column.

The LC-MS method was developed using a Dionex Ultimate 3000RS (ThermoScientifc, CA, USA) module equipped with a binary high-pressure gradient pump, an autosampler,

and a column oven. The column (SeQuant ZIC-cHILIC, 100 x 2.1 mm, 3 μm) equipped with a guard column (SeQuant ZIC-cHILIC, 2 x 2.1, 3 μm) were used for metabolite separation and elution. The mobile phase initial composition, 90% ACN (A) and 10% 100mM ammonium formate (B), was linearly increased to 50% A and 50% B over 15 min, then held for 1 min, followed by equilibration under the initial conditions for another 1 min. The flow rate was 0.3 mL/min and the column was operated at 23±0.1 °C.

A Bruker EVOQ Qube triple quadrupole mass spectrometer (Germany) operated in the positive or negative heated electrospray ionization mode was connected via a divert valve to the LC elution PEEK capillary. The operation parameters were set as the following: Spray voltage +4000/-3500 V, cone temperature 350 °C, cone gas flow 20 psi, heated probe temperature 300 °C, probe gas flow 40 psi (nitrogen), and nebulizer gas flow 45 psi (nitrogen). Argon was used as the collision gas. All other parameters were set at default values provided by the manufacturer. The flow from LC was diverted to waste in the range of 0–1 min and 16.5–25 min. Target metabolites were scanned in the selected reaction monitoring mode (SRM) as fragments of $[M^{+}H]^{+}$ or $[M^{-}H]^{-}$ ions given in Table S4. Metabolite concentrations were normalized to the total metabolite content and relativized to the FGF2-treated cells in 21% $O_2$.
